# Predicting the short-term success of human influenza A variants with machine learning

**DOI:** 10.1101/609248

**Authors:** Maryam Hayati, Priscila Biller, Caroline Colijn

**Affiliations:** Simon Fraser University, Department of Computing Science; Simon Fraser University, Department of Mathematics; Imperial College London, Department of Mathematics

## Abstract

Seasonal influenza viruses are constantly changing, and produce a different set of circulating strains each season. Small genetic changes can accumulate over time and result in antigenically different viruses; this may prevent the body’s immune system from recognizing those viruses. Due to rapid mutations, in particular in the hemagglutinin gene, seasonal influenza vaccines must be updated frequently. This requires choosing strains to include in the updates to maximize the vaccines’ benefits, according to estimates of which strains will be circulating in upcoming seasons. This is a challenging prediction task. In this paper we use longitudinally sampled phylogenetic trees based on hemagglutinin sequences from human influenza viruses, together with counts of epitope site polymorphisms in hemagglutinin, to predict which influenza virus strains are likely to be successful. We extract small groups of taxa (subtrees) and use a suite of features of these subtrees as key inputs to the machine learning tools. Using a range of training and testing strategies, including training on H3N2 and testing on H1N1, we find that successful prediction of future expansion of small subtrees is possible from these data, with accuracies of 0.71-0.85 and a classifier ‘area under the curve’ (AUC) 0.75-0.9.

## 1 Introduction

Human influenza A virus remains a substantial global public health challenge, causing considerable illness and mortality despite the availability of effective vaccines. Influenza viruses are categorized according to features of two surface glycoproteins, hemagglutinin (HA) and neuraminidase (NA), with types such as H3N2 and H1N1 indicating the variant of HA and NA characterizing the strain. Influenza viruses are prone to variability, both in the form of so-called antigenic drift, and in the form of reassortment. Reassortments can give rise to new variants with distinct antigenic properties compared to previous strains; resulting pandemic influenza virus strains may be highly pathogenic. In contrast to pandemic strains arising from reassortment, seasonal influenza virus primarily arises through genetic drift, as influenza virus has a high propensity for generating antigenic variation. This allows influenza viruses to evade host population immunity built up through previous exposure. As a consequence, seasonal influenza virus vaccines need to be regularly updated.

Influenza virus vaccines typically focus on preventing infection by raising antibodies specific to the hemagglutinin (HA) protein. In order to update a seasonal influenza virus vaccine, currently-circulating strains must be selected for inclusion. This relies on surveillance and sequencing of circulating influenza virus genotypes, and on measured antigenic properties of circulating strains. These data do not, in and of themselves, describe future circulating strains, and sometimes the strain selection process does not reflect the near-future composition of the influenza viruses well enough to achieve the desired reductions in illness and mortality. Predictive models are now being used in conjunction with sequencing and immunological surveillance in order to improve the strain selection process.

Phylogenetic trees encode patterns of ancestry and descent in a group of organisms, and so they necessarily include information about differences in these patterns between different subsets of taxa [34, 11]. Trees contain both branch lengths and information in the form of the tree topology or shape. Phylogenetic trees have been used in infectious disease to estimate the basic reproduction number [45], parameters of transmission models [50], aspects of underlying contact networks [38, 22, 39, 28] and in densely sampled datasets even person-to-person transmission events and timing [12, 18, 2, 51]. It is therefore natural to hypothesize that phylogenetic tree structures and branching patterns contain information about short-term growth and fitness. Tree information is central in some predictive models for short-term influenza virus evolution and models of fitness [34, 11]. However, the mapping between the phylogenetic tree structure and interpretable biological information can be subtle, [39, 28, 8, 26] and trees do not directly reveal the short-term evolutionary trajectories of groups of taxa.

Improvements in influenza virus surveillance, sequencing, data sharing and visualization [32, 13] mean that sequence data over considerable time frames is now available to the community alongside intuitive and interactive displays showing how the population of influenza viruses has changed over time. Computational systems to reconstruct large-scale phylogenetic trees from sequence data have also been developed. Machine learning models are well-suited to systematically explore subtle relationships between a suite of features and an outcome. These together present the opportunity to integrate information from different sources to improve short-term influenza virus predictions, using phylogenetic trees as a framework. Here, we use a convolution-like approach to identify small subtrees within a large global H3N2 phylogeny derived from HA sequences sampled between 1980 and July 2018. We fit classification models to detect early signs of growth and hence to predict the short-term success or failure of these subtrees. We validate the predictions on a portion of the data not included in the fit procedure. We relate our predictions to the WHO defined clades [3, 4] for sequences sampled from 2015-2018. Our approach could be performed in real time, is computationally efficient and can be continually refined to improve the quality of predictions as more data are gathered. We suggest that small groups of closely-related influenza virus sequences and the phylogenetic trees that capture their recent shared ancestry patterns can complement other approaches to better predict short-term seasonal influenza virus evolution.

## 2 Results

Briefly, we extract subtrees from the H3N2 phylogeny. Each subtree corresponds to an internal node of the tree and the tip descendants that have occurred within a fixed time frame (1.4 years). The remaining tips occur after the fixed time frame following the relevant internal node, and help to define whether the subtree has successfully grown into the future.

The approach results in a total of 391 subtrees, overlapping to some extent, containing 7615 of the 12785 tips in the full phylogeny. We use a wide range of features of the subtrees, focusing largely on tree structure but also including some branch length features, and the number of changes in the epitope sites of HA compared to previous sequences – see Material and Methods. We train supervised machine learning models to use this information to predict whether subtrees will succeed (see Figure S1). Figure 1 shows the H3N2 hemaglutinin phylogeny and highlights in yellow the tips that belong to at least one subtree.

**Figure 1:**
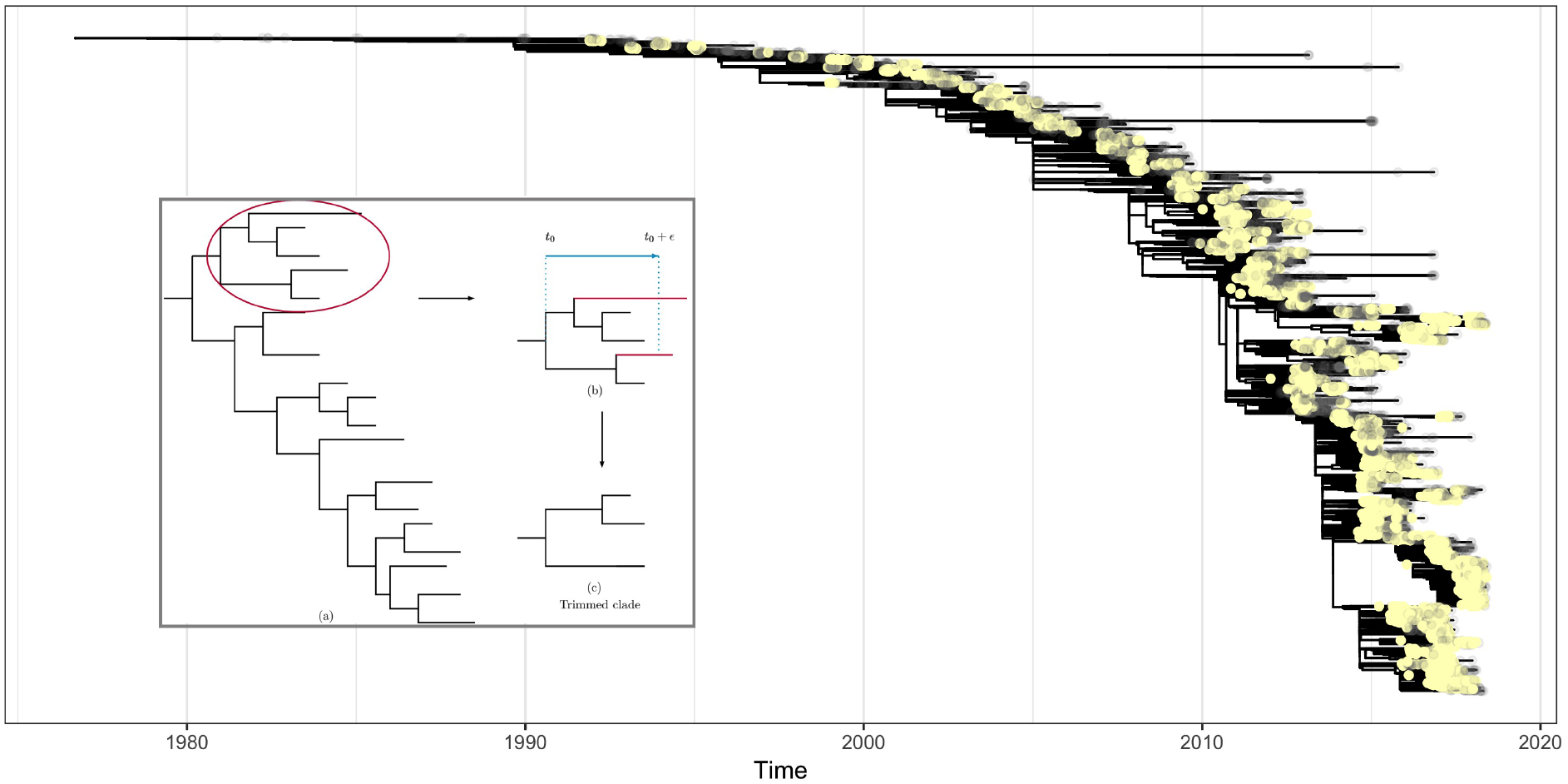
Phylogenetic tree reconstructed from H3N2 subtype sequences using RAxML, with tips highlighted. Each yellow tip is in a trimmed subtree (7615 out of 12785 tips); grey tips are not. The sequences are downloaded from GenBank with dates from 1980 to 2018-5. Long branches in this timed tree did not appear as long branches in the RAxML tree and were not removed (though their tips are not in any trimmed subtree). Inset: illustration of formation of trimmed subtrees: (a) the circled clade contains a subtree; (b) red branches reach tips that occur after the trimming time period and so are pruned out; (c) the resulting trimmed subtree.

The trained models successfully predict which subtrees will grow sufficiently, as measured within 3.4 years of a subtree’s originating node (2 years after the last possible tip in a subtree). Using support vector machine (SVM) classification with a linear kernel, our overall 10-fold cross validation accuracy in H3N2 (using the HA sequences) was 74%. As the accuracy of a classifier can be misleading when there are uneven numbers of samples with the different outcomes, we use the area under the receiver-operator characteristic curve (AUC) to describe the overall performance of our models. The SMV had an average AUC of 0.82 (range 0.73-0.9); see figure S4. We found an accuracy (portion correctly classified) of 79% and AUC of 0.89 when training on 75% of the subtrees chosen uniformly at random, and testing on the rest (Figure 2).

**Figure 2:**
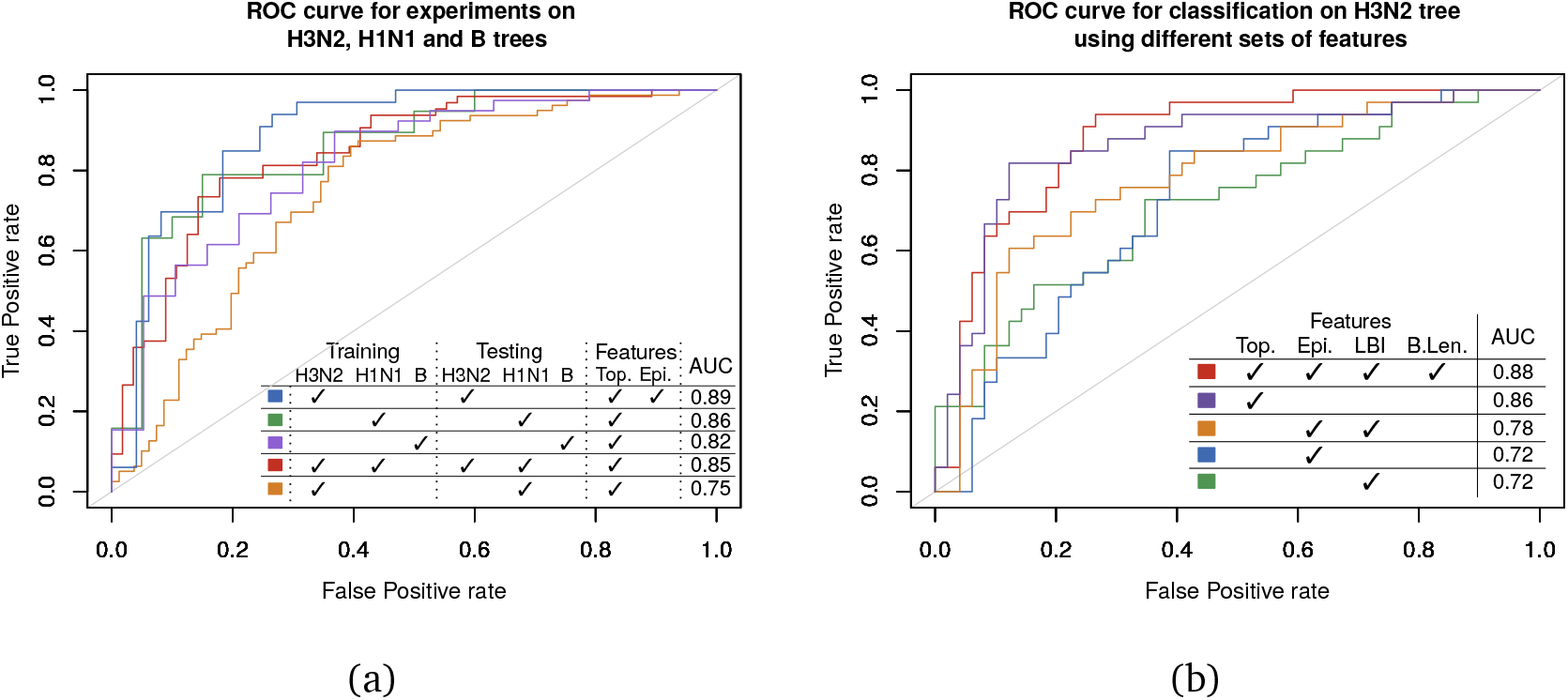
(a) ROC showing performance of linear kernel *SVM* trained and tested on H3N2, trained and tested on H1N1, trained and tested on B, trained and tested on the merged subtrees of H3N2 and H1N1 and trained on H3N2 and tested on H1N1. Figure S4 shows variation in these curves and their AUCs over 10 models trained on 90% of the test data for each case. (b) ROC showing classification performance for restricted sets of features. Top: tree shape features (no branch lengths). Epi: epitope features. LBI: local branching index. B.Len: some features include tree branch lengths.

Figure 2 shows receiver-operator characteristic curves illustrating the trade-off between sensitivity and specificity. AUC ranges were obtained by training 10 models each on 90% of the subtrees; see Figure S4. We obtained a 79% accuracy and 0.86 AUC when we trained a linear kernel SVM model on a training portion of the subtrees (75% of the subtrees chosen uniformly at random) obtained from the H1N1 phylogeny, reconstructed using sequences from 2009 to 2018-05 (see Methods; we did not use epitope features in any H1N1 analyses as the HA protein differs in H1N1). We performed 10-fold cross-validation on H1N1 subtrees, which resulted in 0.76 average AUC (range 0.52-0.95). We also pooled the subtrees of H1N1 and H3N2 and divided the pooled subtrees into training and test data sets; this resulted in an accuracy of 75% and an AUC of 0.85 (10-fold cross-validation AUC range 0.75-0.88). We also applied our method on a phylogenetic tree reconstructed from the HA gene of human influenza virus B. This results in a 0.82 AUC and 78% accuracy when we train our model on 75% of the data chosen randomly and test on the rest. For the influenza virus B tree, we used only the topological features. We also compared classifier performance using only portions of the data, and find that combining the tree shape features with epitope and local branching index (LBI) [34] gives the highest quality, with AUC of 0.88 compared to 0.72 for either epitope features [23] or LBI alone, and 0.78 for these combined.

Our subtrees are based on internal nodes in long-time phylogenies, and these nodes are present as a consequence of the relatedness patterns in all the data that are passed in to the tree reconstruction algorithm (in particular, including sequences from the entire time range). In consequence, a node’s existence and local structure may be conditional on sequences occurring chronologically long after the node. We took several approaches to ensure that our models were not influenced by some such subtle knowledge of the future. We trained models on H3N2 HA phylogeny but tested on an H1N1 HA phylogeny. We obtained an accuracy of 72% and an AUC of 0.75 (range 0.72-0.75 when training 10 models each on 90% of H3N2 subtrees and testing on H1N1; Figures 2 and S4). The reduced accuracy is natural given that the HA proteins differ between the two types. We also created “time slices” from the H3N2 HA sequences using only tips occurring prior to time *i* (*i* ∈ {2012–5, 2013–5, 2014–5,2015–5, 2016–5, 2017–5}). We extracted subtrees, and tested their success using trees reconstructed from all the sequences prior to time *i* + 1 respectively (see Materials and Methods and Supplementary Information). This mimics a “real-time” analysis and ensures that subtrees cannot depend on sequences arising after a set time. This approach performs comparably to our other tests, with accuracy of 0.70% – 76% and an AUC of 0.73 – 0.86.

Subtrees originating after January 2015 (here called “recent subtrees”) did not have enough time to grow into the future, and we do not know whether they are successful, as the predictions are relative to 3.4 years after the initial node. To accommodate for this, throughout our analysis, we only trained and tested models on subtrees that originated prior to January 2015. In this second part, our aim is to make predictions for subtrees whose outcomes are not known. In order to do so, we trained 10 models using 10-fold cross-validation on the non-recent subtrees, as well as a general model, and used these 11 models for predictions on the recent subtrees.

The recent subtrees were labeled according to the clades defined by the World Health Organisation (WHO) (see Figure 3(a) and Material and Methods). Every recent subtree contributed with 11 predictions to its respective clade. The predicted success of a clade in 2019 is the average success of all predictions coming from related recent subtrees. The predictions are presented in figures 3 and S10 (Supplementary Material).

**Figure 3:**
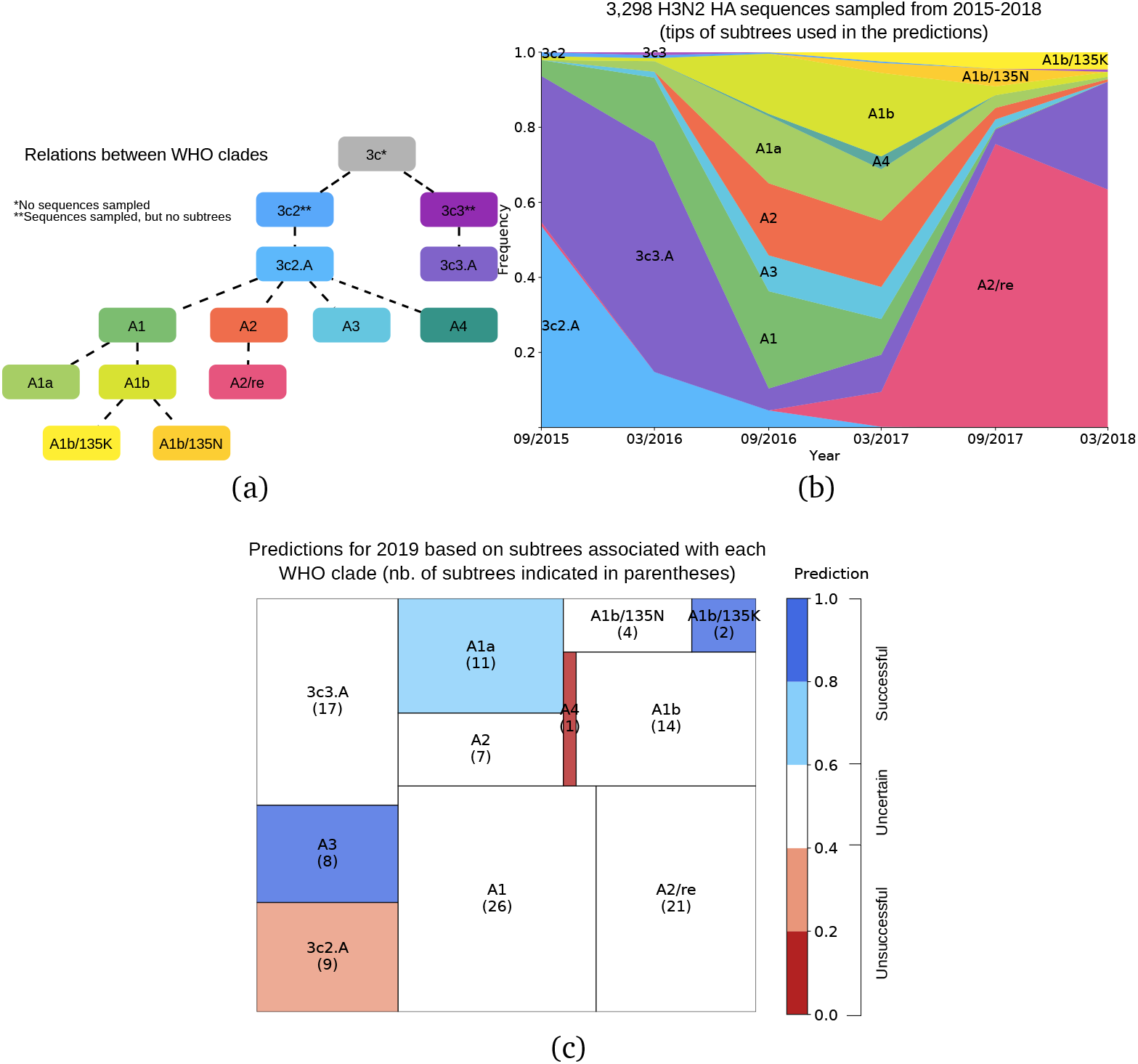
Summary of predictions for the recent subtrees (for detailed information, see Figure S10 in Supplementary Material). (a) Relations between the clades defined by the World Health Organisation (WHO), showing the emergence of new clade designations from existing ones; (b) Frequency of clades through years, based on 3298 H3N2 HA sequences sampled between 2015 and 2018. Every sequence is associated with a tip of one or more subtrees used in the predictions. The clade designation of the tips determines the clade designation of the subtrees by majority rule. (c) Predictions for the recent subtrees. Among the 120 recent subtrees, the outcome (success/failure) of 63 subtrees is not known. Each subtree was tested on 11 different SVM models (see Material and methods). A rectangle corresponds to the prediction of a clade, and its area is proportional to the number of subtrees used in the prediction. The number of subtrees associated with a clade is indicated in parentheses. Colors reflect the combined predictions of the subtrees associated with each clade.

Our recent subtrees originated between 2015-03 and 2017-02, and in this period, contained tips as shown in Figure 3(b), with the majority of tips in clades A1b, A1, A1a, A2 and A2/re. This reflects the sequences in GenBank, and is likely not globally representative [13]. Clades A3, A1 and A1b/135K and A2/re were most strongly predicted to be successful by our measure (fraction of the clade’s subtrees predicted as successful), but in A1b/135K there are only two subtrees on which to base predictions. In clade A3 we predicted 7/8 subtrees as successful, with 5 of those already having shown sufficient growth to meet our success criterion. A2/re has an intermediate signal overall, but has 12 subtrees predicted to be successful. Of these, 10 had already shown sufficient growth to pass our success threshold by the end of our sampling. Indeed the A2/re clade did become very successful, probably due to a re-assortment event [4]. Our model also predicted that other parts of clade A2 (4 of 7 subtrees in A2 but not in A2/re) may grow. In the time frame we had, there were relatively few sequences in our GenBank data that were mapped into the A4, A1b/135K and A1b/135N clades by the ‘augur’ pipeline [13] so these clades have very few subtrees on which to make predictions.

## 3 Discussion

We efficiently predict the success of individual influenza virus subtrees using machine learning tools applied to phylogenetic trees. Our method allows binary classifiers to be trained to predict which currently circulating subtrees will persist into the future based primarily on a suite of phylogenetic features.

Our approach is complementary to previous approaches, including the fitness based model proposed by Łuksza and Lässig [23] and the tree-centred work of Neher and colleagues [34, 33]. Our approach requires a reconstructed timed phylogenetic tree, and can accommodate additional data (e.g. we have used epitope mutations) easily. Other approaches often require additional data such as HI titers and estimates of the ancestral sequences, introducing experimental and computational costs and uncertainty. Our approach makes use of the reconstructed phylogeny in two distinct ways, first in obtaining the groups of taxa (subtrees) considered together for analysis, and second in that the tree shape and length features are derived from the phylogeny, and capture features of the complex branching patterns within subtrees, as opposed to their overall rates.

Our approach is rooted in the hypothesis that fitness and early success leave signatures in the branching time and structure of phylogenetic trees, which can be complemented with additional relevant information such as epitope diversification. With only slight modifications, our model could be applied to other organisms. We could also extend the approach and train regression models to predict the number of tips arising from subtrees. However, a natural limitation of this (and other tree-based approaches) is that it detects signs of early growth - if an adaptive new mutation arose in the population and was sampled before that early growth could occur and be sampled, then we could not detect early signs of growth and would not see the new adaptive mutation. In contrast, a principled modelling approach based on an understanding of both what makes an influenza virus fit and on the current composition of population immunity would likely be able to detect fit novel mutations without relying on such viruses already having begun to spread.

Influenza virus A can be categorized based on the presence of different proteins on the surface of the viruses: hemagglutinin (HA) and neuraminidase (NA). We use trees reconstructed from HA sequences; relatedness in the HA tree corresponds to similar HA sequences and hence to similar immunity profiles, as antibodies are induced by HA. Indeed, path lengths in the HA phylogenetic tree provide a good model for antigenic differences modeled by serological assays [34]. Trees describe the relative number of recent descendants of a lineage compared to closely-related lineages, the timing and asymmetry in the descent patterns and the short- and long-term future populations that are related to the lineages. Our approach allows this information to be included in predictive efforts.

We did not include information about proximity of strains to current or recent vaccines, which might have led to false positives in our results, if a subtree showed early signs of success but was later suppressed by vaccination (this seems unlikely, as only a small portion of the global population are vaccinated). We also did not explicitly include immunological assay data, as these are not generally available. We do not have good estimates of the current frequencies of subtrees or strains - indeed, if up-to-date global frequencies were available at high resolution it would greatly facilitate short-term prediction. We used epitope sites following the approach of [23]; a model reflecting the impact of polymorphisms across more locations in HA and in other genes, if this were available, would also potentially improve predictions.

We used RAxML to infer the trees; it uses a maximum likelihood approach and is considered a state-of-the art reconstruction algorithm [52, 20]. Due to the large numbers of isolates we did not perform Bayesian Markov Chain Monte Carlo (MCMC) tree reconstruction to accommodate tree uncertainty; in addition to each required MCMC run, this would also have required exploration of different priors and assumptions, and it is computationally unfeasible for thousands of tips. In order to check the robustness of our approach, we used different training and testing trees including training on H3N2 but testing on H1N1, pooling H3N2 and H1N1 and using distinct time slices and consistently obtained successful predictions.

Tree shape statistics are dependent on the size of a subtree, so in order to have a more robust comparison between the subtrees, it would be best to select subtrees with approximately the same size. In our data the size of a trimmed subtree was a poor predictor of the fractional growth. Furthermore, constraining the sizes of subtrees reduces both the number of subtrees and the number of tips that can be included in the analysis, and size may in fact be a valuable predictor. We chose the approach here to balance these contrasting issues; the ongoing sampling and sequencing and the natural passing of time will ultimately provide more data – more years, and more samples per year – such that subtrees of more consistent size can be used. We anticipate that this will increase the quality of predictions. Another next logical step will be to model competition or other interactions between major lineages. We have not explicitly modelled ancestral states, key individual mutations, serological data or estimates of the fitness of sequences, but our approach could easily integrate results from models that include these features. Ideally, all relevant sources of information would be integrated and updated in real time [33, 23]. However, while short-term forecasting based on various data sources is feasible and is required to update seasonal vaccines, perfect short-term prediction and accurate long-term prediction are likely not possible because evolutionary events are fundamentally stochastic.

Recent studies on the viral isolates from vaccinated individuals indicate that they are significantly distinct from the vaccine strain and are broadly distributed on the tree, resulting in accelerated antigenic evolution [47]. Researchers have been working to develop a universal vaccine that would provide broad protection against both seasonal and pandemic influenza. Recent studies have also indicate that universal vaccines could decelerate the speed of evolution [31, 47]. If successful, such universal vaccines would eliminate the need for continual updates to seasonal influenza virus vaccines, but we would suggest that even so, current efforts to make short-term predictions based on surveillance and sequence data gathered over time can yield both practical results and broader insights into short-term patterns of evolution. Our approach indicates that fitness can affect the phylogenetic properties of a tree reconstructed from influenza viruses and that properties of small subtrees can be used as a set of predictors to estimate which groups of sequences are showing signs of success.

## 4 Methods and Materials

Our approach is rooted in the hypothesis that fitness – the reproductive rate and capacity of a group of organisms – affects tree structure and branching patterns (including timing) and that this information can be extracted using machine learning tools.

### 4.1 Definitions

Given a phylogenetic tree *T*, a tip (also called an external node) of *T* is a node of degree one. An internal node of *T* is any non-leaf node of the tree. A rooted tree is a tree in which a particular internal node, called the root, is distinguished from the others; it is usually considered to be a common ancestor of all the other nodes in the tree. In a rooted tree *T*, the parent of a node *i* is the node preceding it on the unique path from the node to the root *r* of *T*; all nodes of *T* except its root have a parent. A child of a node *i* is a node whose parent is *i*. A phylogenetic tree is bifurcating if all its internal nodes have two children. We use a rooted bifurcating phylogenetic tree that is reconstructed from the hemagglutinin sequences of influenza virus A strains (by maximum likelihood).

### 4.2 Reconstructing Influenza virus trees

We collected full-length HA sequences from human H3N2, H1N1 and influenza virus B on GenBank [5, 43]. We used unique sequences of H3N2-HA from human cases for years between 1980 and 2018-5, excluding laboratory strains. This results in approximately 12919 sequences. For influenza virus H1N1 we collected pandemic sequences from 2009 until 2018-5, giving 10652 unique sequences. For influenza virus B, we used sequences from 1980 to 2018-5, resulting in 7257 unique sequences. We aligned each set of sequences using MAFFT [14, 15] and then we used RAxML [46] to reconstruct maximum-likelihood phylogenetic trees. The reconstructed trees using RAxML are neither rooted nor dated; in some cases they include very long branches, which we removed. We rooted trees with the *rtt* function in the *ape* package [37] in *R* and then converted them to timed trees using the Least Squares Dating (*LSD*) software [48].

### 4.3 Subtree extraction

We use a convolution-style approach to identify subtrees of the global timed phylogeny that serve as units of analysis. For each internal node i in the tree, we find the tips that occur within a fixed time window (1.4 years by default) chronologically following *i*; this is node *i*’s “trimmed subtree”. We cannot train machine learning models on the subtree descending from every internal node in the tree, because these subtrees will overlap substantially. We use the notion of a node’s “relevant ancestor” (described below) to control the overlap, and select subtrees in a convolution-like way.

We first initialize each node’s relevant ancestor to be its parent. We traverse the nodes of the tree in a depth-first search order. If a node’s complete subtree is too small (fewer than 8 nodes by default), we reject the node and all its descendants, as none of the descending nodes can have a larger subtree than the node itself. If the node’s trimmed subtree is too small but its complete subtree is large enough, we reject the node but not its descendants, since they may have subtrees that are large enough. If the node’s trimmed subtree is large enough, we check the overlap between the node’s subtree and its relevant ancestor’s subtree. The overlap is the portion of node *i*’s trimmed subtree that is contained in the relevant ancestor’s trimmed subtree. If this overlap is not too large (under 80% of the subtree size by default), the subtree is included in our analysis. If the overlap is too large, we reject the node, and we set the relevant ancestor of the node’s children to be the node’s relevant ancestor. In that way, when we decide whether to accept the subtree of the node’s child, we will control the correct overlap.

In this way, we obtain subtrees containing tips that are within a specified time window after their originating node, have at least a minimum number of tips, and have a limited overlap with other subtrees. We varied the minimum size, time interval and permitted overlap (See table S1 and Supporting Information). We obtain a total of 392 subtrees in H3N2, and 198 subtrees in H1N1. After removing subtrees with big size in relation to the average size, and recent subtrees with insufficient growth to determine their outcome, we obtained 329 subtrees for H3N2 and 160 subtrees for H1N1.

### 4.4 Features

We use a set of features defined on subtrees, including both tree shape and patterns in the branch lengths. The topological features are summarized in Table 1. For the H3N2-HA dataset, we also consider some features derived from the epitope sites of the tips of the subtree. For each subtree, we consider the mean, median and the maximum genetic distances between the epitope sites of the tips of a subtree and the epitope sites of the sequences with dates prior to the subtree. We used the locations of known antigenic epitopes as mentioned in [44], namely 72 sites in the *HA1* subunit of HA.

**Table 1:**
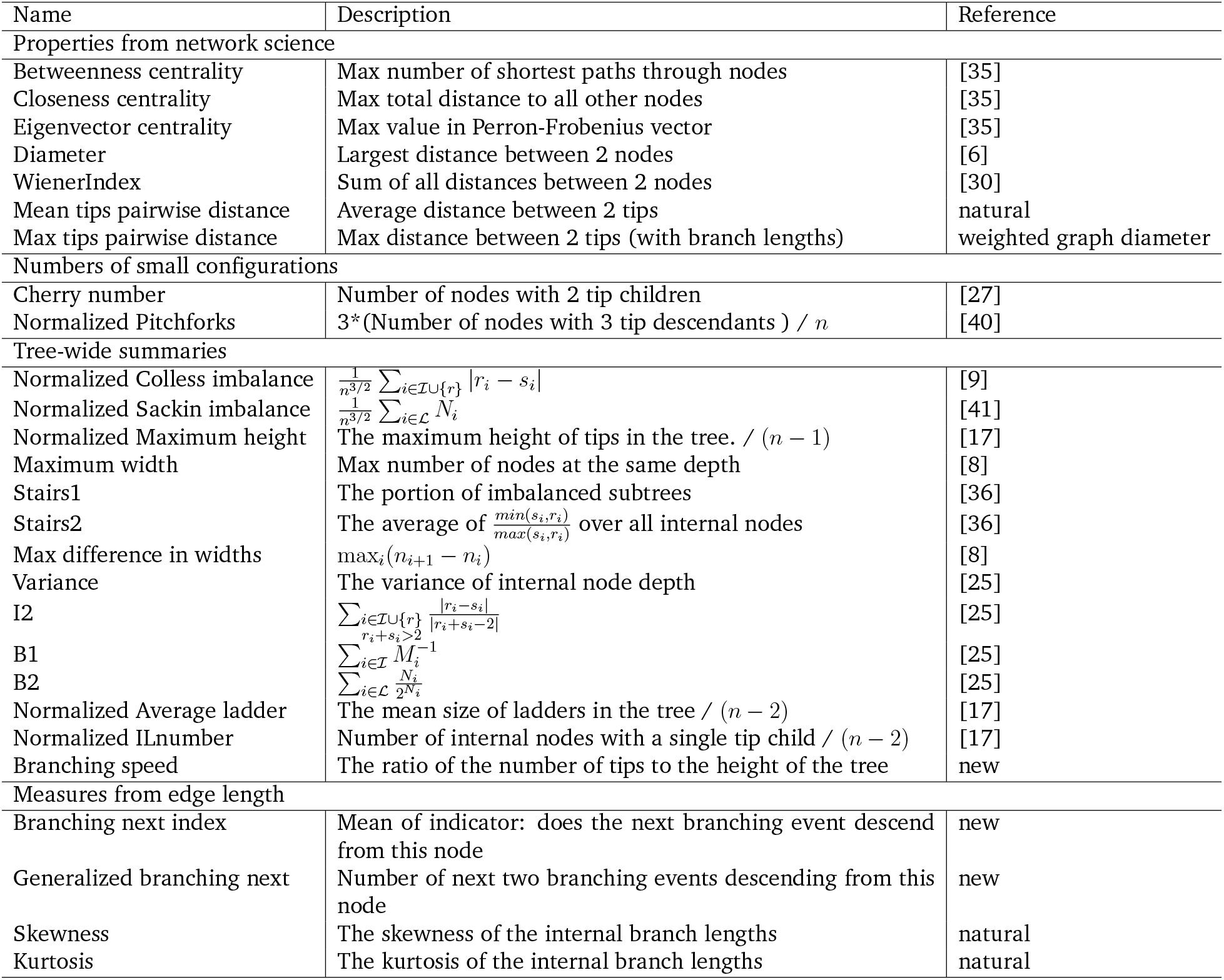
Brief definition for tree shape statistics. Here *r_i_* and *s_i_* are the number of tips of the left and right subtrees of an internal node respectively, *n* is the number of tips of a subtree. *n_i_* is the number of nodes at depth *i, M_i_* represents the height of the subtree rooted at an internal node *i*, and *N_i_* is equal to the depth of node *i*. A ladder in a tree is a set of consecutive nodes with one tip child. We represent the set of all internal nodes of a tree by 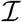, the set of all tips (or external nodes) by 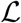, and the root of a subtree by *r*. In “generalized branching next” we chose *m* = 2. Skewness and Kurtosis are two measures to describe the degree of asymmetry of a distribution [24]. The tree shape statistics induced by betweenness centrality, closeness centrality and eigenvector centrality are defined as the maximum values of each centrality over all the nodes of a tree, and distances are in units of number of edges (without branch lengths). Features called “natural” may not have been used as tree features previously but are natural extensions of simple features (eg skewness is a natural quantity to compute). The network science properties were computed in R using the treeCentrality package [21] and the tree-wide summaries were primarily obtained using the phyloTop package [1].

Our features cover a wide range of global and local structures in trees, expanding considerably over previous approaches which largely focus on tree asymmetry and a few properties of branch lengths [34, 11]. Previous authors have noted that fitness leaves traces in genealogical trees [11] by observing in fixed-size populations that increased fitness resulted in increased asymmetric branching and long terminal branch lengths; Neher and colleagues used the local branching index (LBI), a measure of the total branch length surrounding a node, in their predictive model [34]. We significantly expand on the repertoire of tree features, including asymmetry and measures of local branching but also including features derived from network science that capture global structure of the subtrees, small shape frequencies and others – see Table 1.

For comparison purposes we implemented Neher et al.’s local branching index (LBI) [34], which is a measure of rapid branching near a node in the tree. In doing this we noted that there are strong parallels between the LBI and the weighted version of the Katz centrality, a classic measure from network science [16]. Figure S8 shows the correspondence. We performed the main classification task (H3N2) using all our features, only the topological tree features, only the epitope and LBI, only the epitopes and only the LBI (Figure 2(b)). We found that the combined features gave the best performance, followed by the tree features.

### 4.5 Success and training approach

We call a subtree of size *n* “successful” if its root has a total of more than *αn* tip descendants in the time frame of 3.4 years from the root of the subtree. The threshold of *α* = 1.1 results in a good balance of successful and unsuccessful subtrees, which facilitates training the machine learning models (see Supporting Information). We chose to use fractional growth as our outcome rather than proximity to tips of the following season, because proximity to the following season fluctuates depending on when in the season the subtree originates, the definition of the season (i.e. the cutoff dates) and the subtree’s location (tropical vs temperate).

The potential overlap between subtrees could induce dependence in the outcome variable (success), i.e., if nodes *n_i_* and *n_j_* have overlap, and *n_i_* is successful, then *n_j_* may be more likely to be successful. Notice that having some tips in common does not mean that overall subtree features are similar, but we hypothesize that the chance increases as the proportion of shared tips grows. If *n_i_* was in the training data and *n_j_* in the test data, and if the two subtrees had similar features, then correlations in the success of these two subtrees could result in overfitting the data. Controlling this potential effect is one reason to use a cutoff of 3.4 years for success (meaning that *n_j_* could be unsuccessful with *n_i_* successful). In one of our experiments, we trained on H3N2 and tested on H1N1 (see Figure 2(a)) to ensure that test and training data are completely distinct. We also explored the performance of our approach on a set of pool subtrees from both H1N1 and H3N2.

Our data are censored, because we cannot observe the future of subtrees beginning in 2018; furthermore, we have limited knowledge of the true success of subtrees beginning in the most recent 3.4 years of our data (since it takes approximately 2 years following the end of the trimmed subtree before its success is known). We only know whether a subtree has been successful if it has already had a sufficient number of tips; other subtrees may yet do so. Accordingly, we could not train our models using the last few years of data. We used 10-fold cross validation on the set of subtrees whose ancestral nodes arose before 2015-1. We refer to this as the set of “non-recent subtrees”. This results in one “out of fold” prediction for each such subtree. We trained an additional “general” model on the set of non-recent subtrees. We then had 10 cross-validation models and one additional model that we could use to make predictions on the subtrees arising after 2015 (for which the true success is only partially known).

The structure (and hence internal nodes) of a large phylogeny depends on all tips, not only the tips prior to the node chronologically. To avoid having the “future” tips affect the nodes on which we based our analysis, we also used a “time slicing” approach that is amenable to use in real-time, season by season. Here, we extracted subtrees not from the full phylogeny but from a tree containing only tips prior to a fixed time. We then assessed the success of these subtrees with reference to later tips (see Supporting Information).

### 4.6 Classification

We use several different binary classification tools, including support vector machine (SVM) with a range of kernel choices [10]. We use *R* implementations in the packages *e1071* [29]. For all the experiments, we randomly chose 75% of our subtrees for training the model and left the rest for testing. We selected a training and a test dataset (75% for training and the rest for testing) and perform 10-fold cross validation on the training dataset alone; this is to find the best gamma and cost parameters without using all the data to do so, see Figure S1. Among different binary classification tools that we used, SVM with a linear kernel had the best performance on this 75% training set, so we proceeded with this option for the remaining results. Datasets can have outliers that affect the training process. We use the local outlier factor (LOF) algorithm [7] implemented in the *DMwR* package [49] in *R*. For classification on the H3N2 tree, we removed 5 outliers which mostly were large subtrees, most of whose descending tips were contained in other subtrees. In the H1N1 tree, we had 198 subtrees; we removed the largest 11 as outliers, for the classification. For the experiment on the merged H1N1 and H3N2 subtrees we removed large subtrees of both H3N2 and H1N1 to have a set of subtrees of approximately the same size. In the experiments on the influenza virus B tree and the combinations of influenza virus B and other types, we remove large subtrees to obtain a set of trees of approximately the same size in the training and testing groups. Our training and testing scheme is illustrated in Figure S1.

### 4.7 Clade assignment

We implemented a Python script using the same approach as the one found in *augur* (https://bedford.io/projects/nextflu/augur/) to assign a clade to a sequence. A clade is defined by a set of amino-acid or nucleotide mutations specific to that sublineage. All mutations used as markers occur in the HA1 and HA2 proteins of H3N2-HA sequences, and the detailed list of mutations can be found at https://github.com/nextstrain/seasonal-flu/blob/master/config/clades_h3n2_ha.tsv. A sequence is assigned to a clade only if it satisfies all criteria of the clade, i.e., only if the sequence contains all the specific mutations.

It is possible for a sequence to satisfy the criteria of more than one clade, since some clades are descendants of previous ones. For instance, the sequence KU289613_A/Corsica/33-02/2015_2015/02/20 satisfies requirements for clades 3c, 3c3 and 3c3.B. In this scenario, the most recent clade is assigned (3c3.B). Among the 12919 sequences analyzed, 1106 sequences were assigned to exactly one clade, 7487 sequences were assigned to more than one clade, and 4326 sequences had no clade assigned. Most of the unassigned cases correspond to older sequences (Figure S7). Among the sequences assigned to more than one clade, we used the clade relations presented in Figure 3(a) to determine the most recent clade (e.g. A1a is contained within 3c2.A, though this information is not encoded in the clade names). The relations between clades were manually inferred from the tree at https://nextstrain.org/flu/seasonal/h3n2/ha/3y.

The clade designation of the tips determines the clade designation of the subtrees by majority rule. As a result, in the predictions presented in Figure 3, some older clades (3c2 and 3c3) have tips assigned to them (9 and 4 tips, respectively) but no subtrees and, therefore, no predictions.

## Acknowledgments

CC and PB would like to acknowledge the Canada 150 Research Chair program. CC was also funded in part by the Engineering and Physical Sciences Research Council of the United Kingdom (EP/K026003/1). MH would like to thank Dr. Leonid Chindelevitch for providing this visiting scholar opportunity through funding from an NSERC Discovery Grant as well as the CANSSI Collaborative Research Team.

## 5 Supplement

### Training, Model parameters and results

We have used cross-fold validation to find hyperparameters (eg SVM with a linear kernel) that perform well in the classification tasks. To avoid inflating estimates of the accuracy of the resulting classification, we chose hyperparameters based only on the training data. Our training approach is summarised in Figure S1.

**Figure S1:**
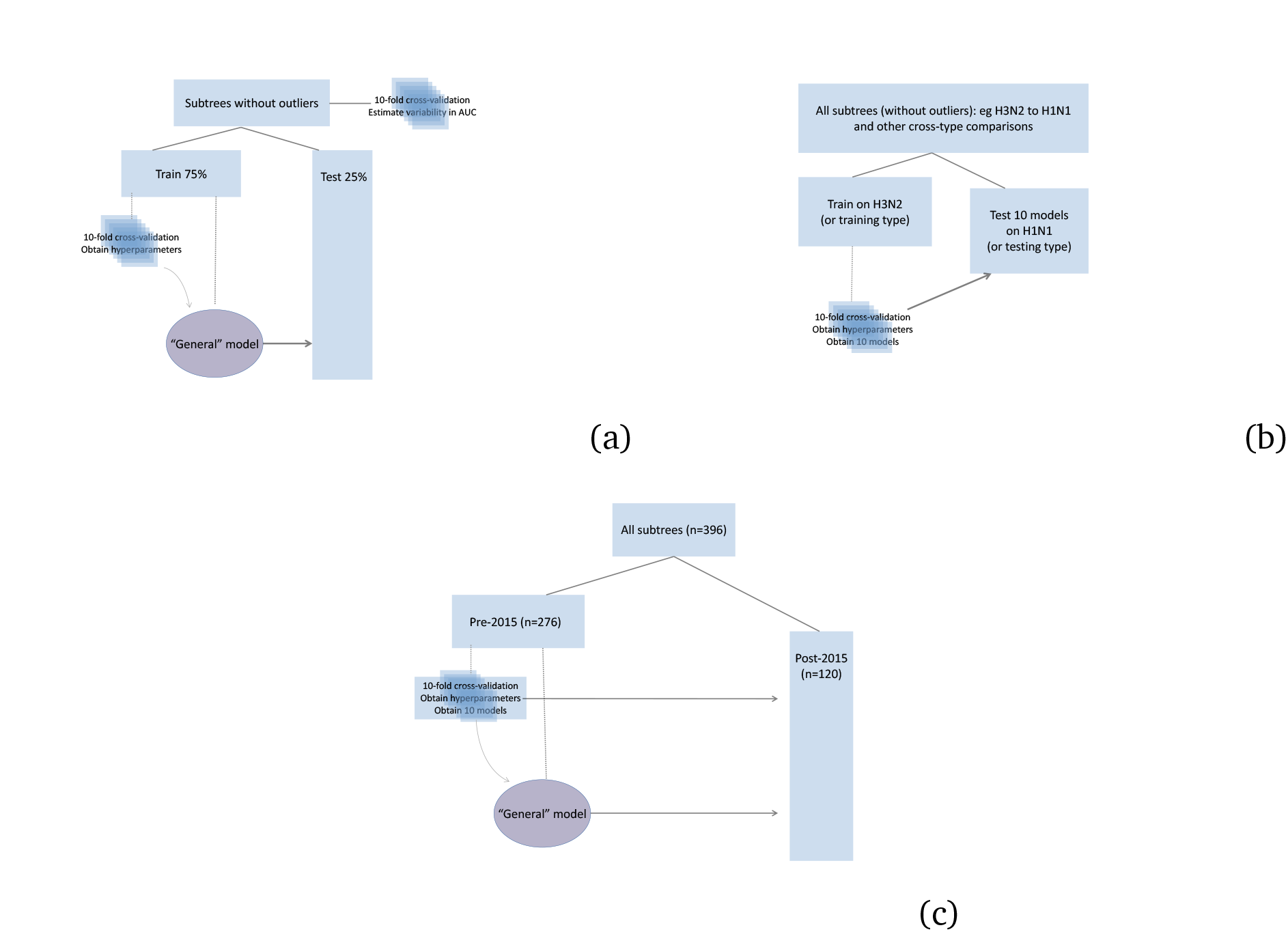
(a)This figure illustrates our training and testing approach for experiments on influenza virus H3N2, influenza virus H1N1, influenza virus B and pooling the subtreees. We divide the subtrees whose outcome is known into training (75%) and testing (25%) data and choose hyperparameters using 10-fold cross-validation on the training data only. We use those hyperparameters to train the “general” model and test it on the training data. (b) The schematic figure for training on a tree (H3N2) and testing on another tree (H1N1).(c) For the prediction task, we divide our subtrees into recent (subtrees originating after 2015-1) and non-recent sets (the rest of the subtrees). We then trained 10 models using 10-fold cross validation together with one general model on the set of non-recent subtrees and used these 11 models for prediction on the recent subtrees.

Because we have used a fixed time frame to select subtrees, the subtrees vary considerably in size (since we do not control size). Successful subtrees are larger (median 18) than unsuccessful ones (median 12). Figure S2 shows the sizes of the trimmed subtrees from H3N2 that were used in the main analysis, with bars shaded to the outcome. We observed that size alone does not successfully classify the success of subtrees, though it contains some of the information (AUC 0.63). We chose to control time frame rather than subtree size, as time frame has a clear biological meaning and the size of subtrees may in fact be useful information for the classification; trees with rapid branching can achieve more tips in the fixed time frame. However, size (and apparent rapid branching) may also reflect sampling differences.

**Figure S2:**
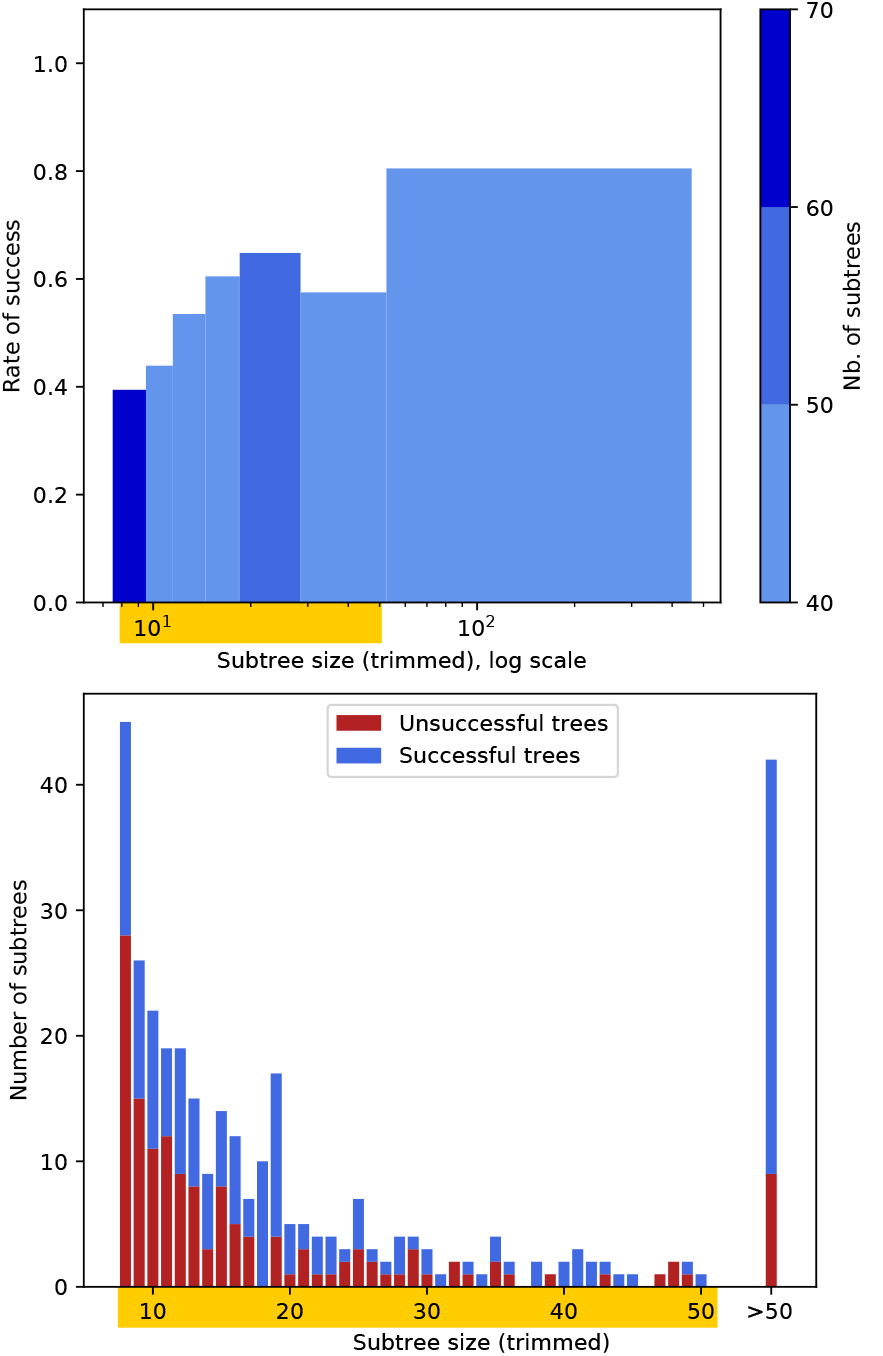
(Top) Correlation between the size of trimmed subtrees (x-axis) and the rate of success (y-axis) for the 391 subtrees from the H3N2-HA dataset. The rate of success is defined as the number of succesful trees divided by the total number of trees for a certain range of sizes. The ranges were computed in order to encompass approximately the same number of subtrees, and the color of the bars represent how many subtrees were taken into account for the computation of the success rate. The subtrees vary from size 8 up to 460, but as most of the dataset is composed by small subtrees, we used a log scale to better visualize the information. The subtree sizes highlighted in yellow are detailed in the bottom panel. (Bottom) Subtree size distribution for trees up to size 50, which corresponds to 87.4% of the dataset. For each size, the graph shows how many subtrees were succesful (blue blar) and unsuccesful (red bar).

We used three sets of parameters to extract the subtrees of influenza virus H3N2. These three models are summarized in Table S1, and the results of prediction using each of these models are shown in Figure S3.

We also explored changing the success threshold, defining a subtree of size *n* as successful if its ancestor eventually has at least one more tip, more than 1.1*n* tips and more than 1.2n tips (ie if size > *αn* for *α* =1,1.1,1.2). Figure S3 shows the performance under these variations.

**Table S1:** Our subtree selection algorithm uses three parameters: a minimum subtree size, a maximum allowable overlap and the length of the time window (Figure 1). We denote our default as “ALT0” and our alternatives as “ALT1” and “ALT2”. There are natural trade-offs: a larger minimum size, lower overlap and longer time frame all result in fewer accepted subtrees. We found good performance for each of these three alternate setups.

| Model | MinTotalSize | MinTrimSize | OverlapCutoff | TimeFrame |
| --- | --- | --- | --- | --- |
| ALT0 | 8 | 8 | 0.8 | 1.4 |
| ALT1 | 12 | 12 | 0.95 | 2 |
| ALT2 | 7 | 7 | 0.7 | 1 |

**Figure S3:**
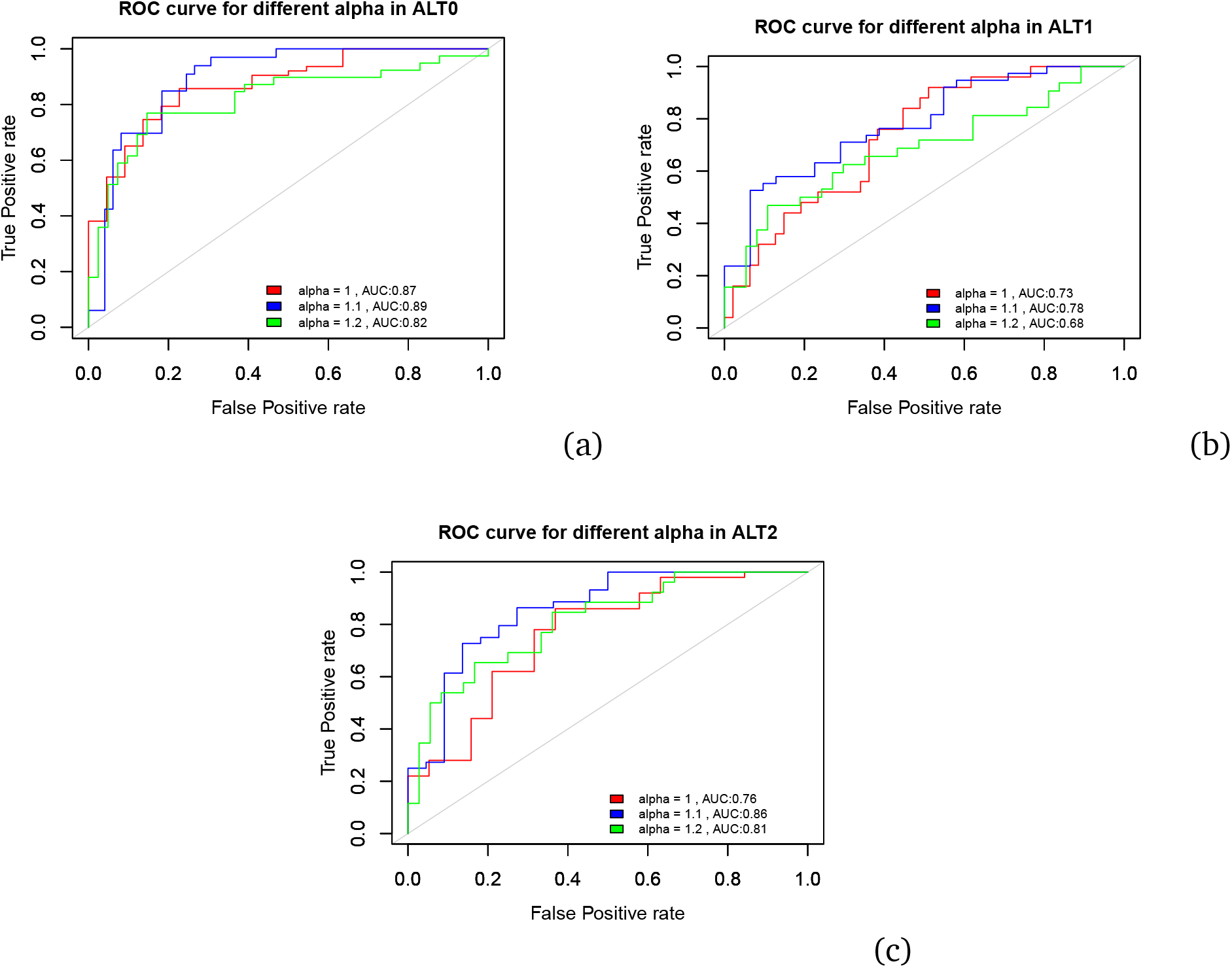
The ROC curves from prediction for the H3N2-HA dataset using different models and different definitions of a successful subtree. In all models, setting the threshold to an appropriate value allows a balance between successful and unsuccessful outcomes, resulting in better performance. Among different sets of parameters for extracting the subtrees, ALT0 is the most powerful model resulting in 0.89 AUC. In all of these experiments we used the topological and epitope features.

### Cross-validation on different experiments

To determine how variable the AUC and accuracy figures are for the analyses reported in the main text, we performed 10-fold cross validation for the main prediction on H3N2, the analysis on H1N1 only (trained on a portion of the H1N1 subtrees), the pooled analysis of H3N2 and H1N1 and the predictions on H1N1 using a model trained on H3N2. The resulting ROC curves are shown in Figure S4, (a)-(d) respectively. AUC values are consistently above the random classifier’s expected value of 0.5, with ranges given in the caption of Figure S4. It is particularly encouraging that the test on H1N1, using a model trained on H3N2 (Figure S4(d)) has a highly consistent set of AUC and accuracy values (range of AUC 0.75-0.88) because this test is in some sense the most challenging; no subtree in the test data (H1N1) ever shares a tip with a subtree in the training data (all H3N2) and the risk of overfitting is low (also note that we did not select a model or method based on this test). The same holds for models trained on H3N2 and tested on influenza virus B, and so on. The most variable set of AUCs arises from H1N1 alone, which is likely due to the lower volume of data.

**Figure S4:**
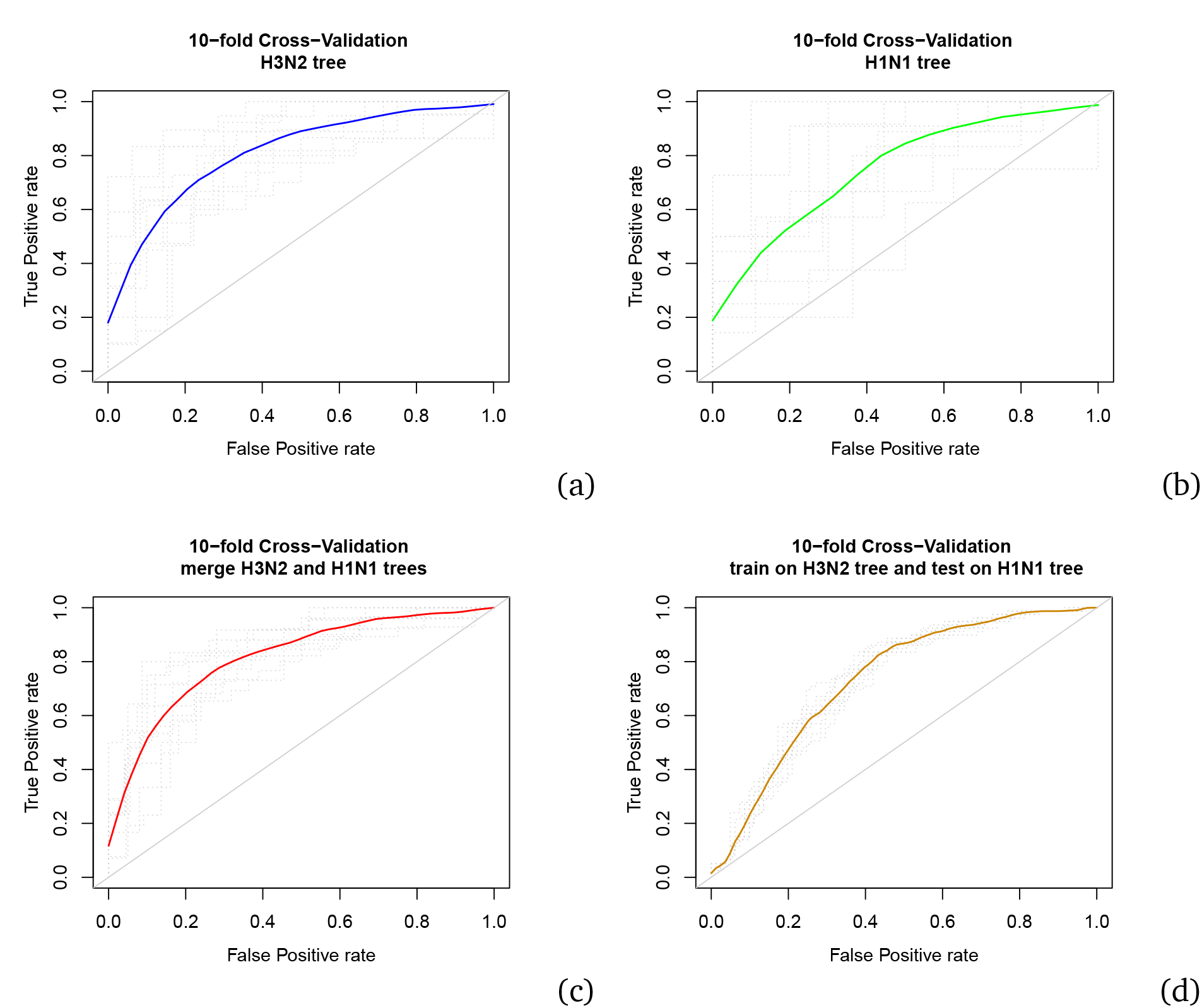
(a) The result of 10-fold cross validation for *SVM* models trained on the subtrees whose ancestral nodes occurred pre-2015-1 or which began later but whose outcomes are known. AUC values range from 0.73 to 0.90 (average 0.82) with 80% of the folds resulting in more than 0.75 AUC. (b) 10-fold cross-validation on the H1N1 tree using *SVM* with linear kernel and a set of topological properties of the clades. AUC values range from 0.52 to 0.95 (average 0.76) with AUC more than 0.75 in 70% of the folds. (c) The result of 10-fold cross-validation on the merged dataset of H3N2 and H1N1 trees. The minimum and maximum values of AUCs among the folds are 0.75 and 0.88 respectively (average 0.82). (d) 10-fold cross-validation for training on the H3N2 tree and testing on the H1N1 tree; we removed 10% of the training data at each fold and evaluated the model on the test data. AUC values range from 0.72 to 0.75 (average 0.74).

### Time slices

To ensure that the method does not rely on internal nodes whose existence depends on the full dataset, we reconstructed the influenza virus H3N2 tree using only the sequences observed prior to time *i* (*i* ∈ {2012 – 5, 2013 – 5, 2014 – 5, 2015 – 5, 2016 – 5,2017 – 5}); call this tree *T_i_*. We extracted the subtrees of this tree using the ALT0 parameters. Naturally, the T_i_ tree does not contain the information as to whether its later-occuring subtrees grow into the following season. To find the remaining success information, we used the tree reconstructed from the sequences any time up to *i* + 1 (*T*_*i*+1_). Consider a subtree *c* in the *T_i_*, with tip set *S_c_* and size *n*. First we find the most recent common ancestor (*MRCA*) of *S_c_* in the *T*_*i*+1_. Then, we compare the size of the subtree *c*(*n*) with the size of the subtree rooted at the MRCA of *S_c_* in *T*_*i*+1_ (*m*). If *m* > *αn* (*α* ∈ {1.1,1.2,1.3,1.5}) then we say that subtree *c* is successful (see Figure S5). Again we tried different *α* cutoffs to obtain a balanced dataset. (see Figure S6). We randomly choose 75% of the subtrees for training our model and leave the rest to test the model. In order to find the hyper-parameters of the model we performed 10 fold cross-validation on the training set. We tried *linear, radial* and *polynomial* kernels for the SVM and, among these, the linear kernel resulted in the best performance (data not shown).

**Figure S5:**
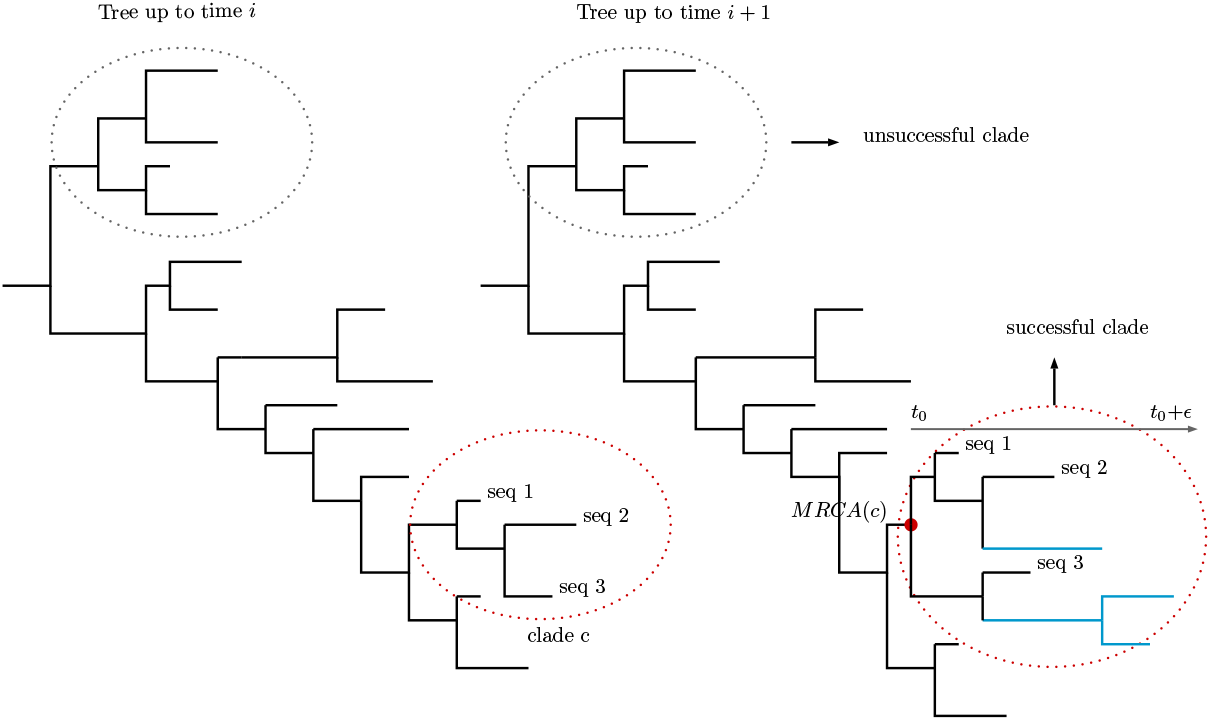
This figure depicts the definition of a successful subtree in the time slicing approach. The tree on the left, *T_i_*, represents a tree reconstructed from a set of sequences up to time *i* and the tree on the right shows the same tree after one year (*T*_*i*+1_). We compare these two trees to predict the successful subtrees in the time slicing approach. For each subtree *c* we compare the size of the subtree in *T_i_*(*X*) with the size of the subtree rooted at the most recent common ancestor of the tips of subtree *c* in tree *T*_*i*+1_(*Y*), in an interval of 3.4 years following the root of the subtree. If *Y* > *αX* (*α* ∈ {1.1,1.2,1.3,1.5}) then we say subtree *c* is successful. This overcomes the challenge that *T*_*i*+1_ does not contain the root of *c*; it does contain a node that is the MRCA of the tips in subtree *c*.

**Figure S6:**
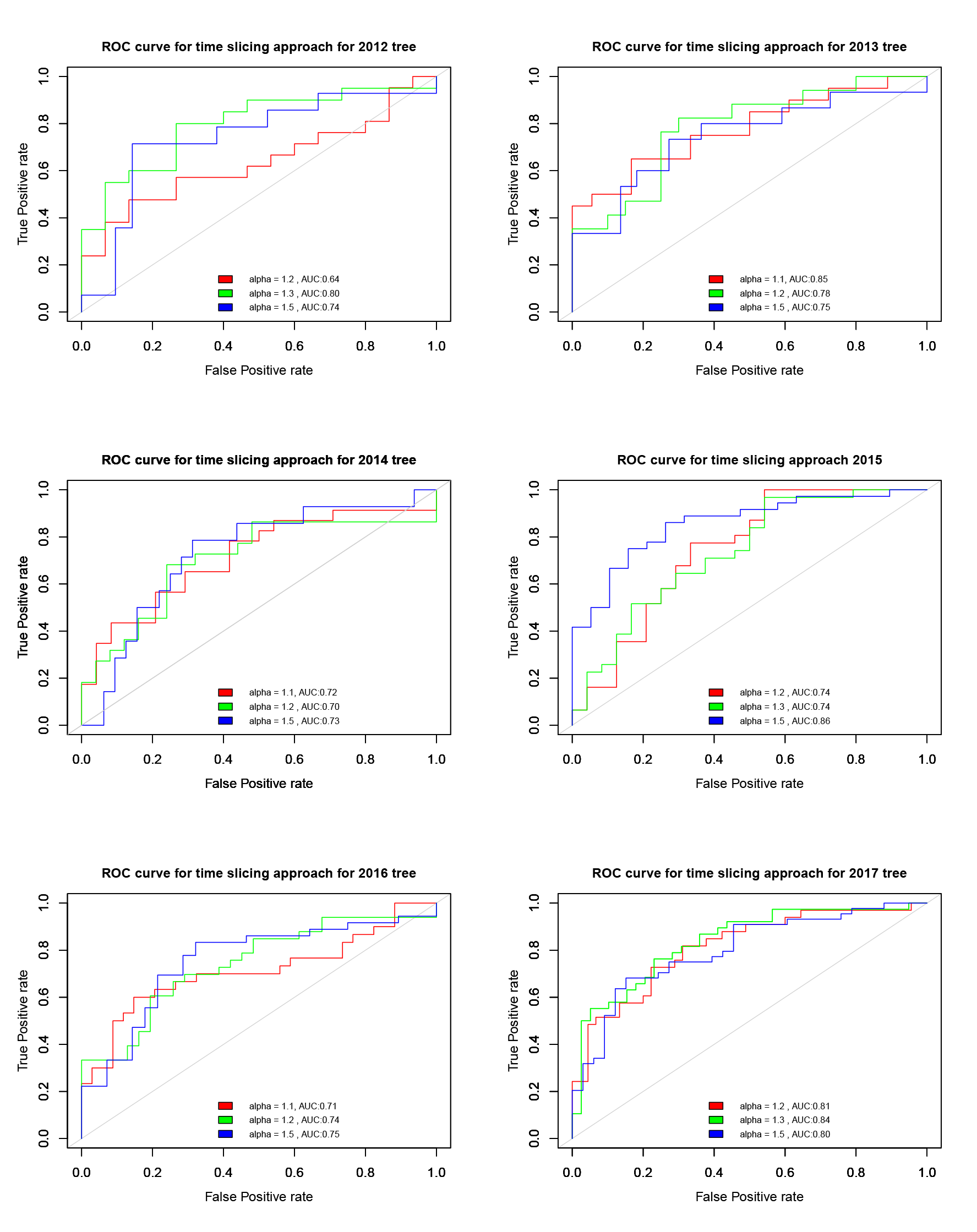
We classified the subtrees of the influenza virus H3N2 tree reconstructed from the sequences up to time *i* by comparing with the tree reconstructed from sequences up to time *i* + 1 which *i* ∈ {2012 – 5, 2013 – 5, 2014 – 5, 2015 – 5, 2016 – 5, 2017 – 5}. In this figure the result of classification using different ratio between the size of a subtree in time *i* and the size of the corresponding subtree in time *i* + 1 is shown.

**Figure S7:**
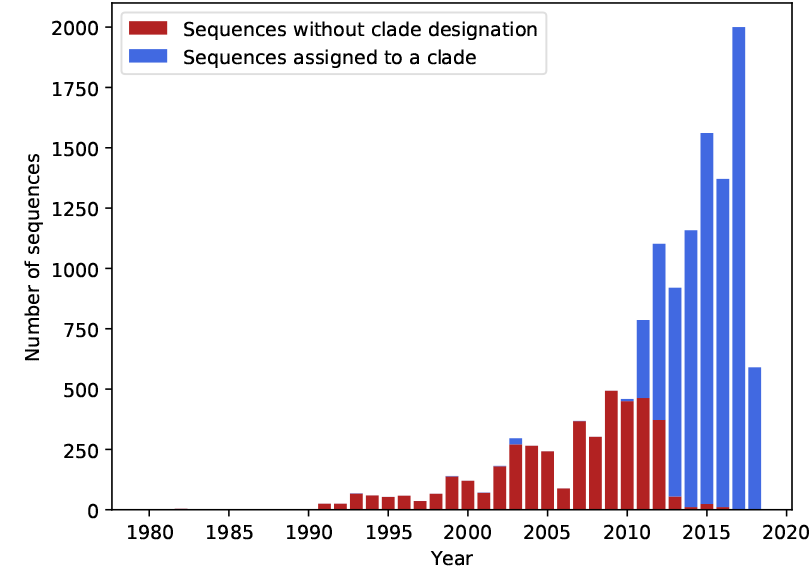
Distribution of H3N2-HA sequences between 1980 and 2018. The blue bar corresponds to sequences assigned to one or more clades, whereas the red bar corresponds to sequences without clade designation. The rate of unassigned cases drops considerably for recent sequences (from 2011 on), which encompass most of the analyzed dataset (73.45%).

**Figure S8:**
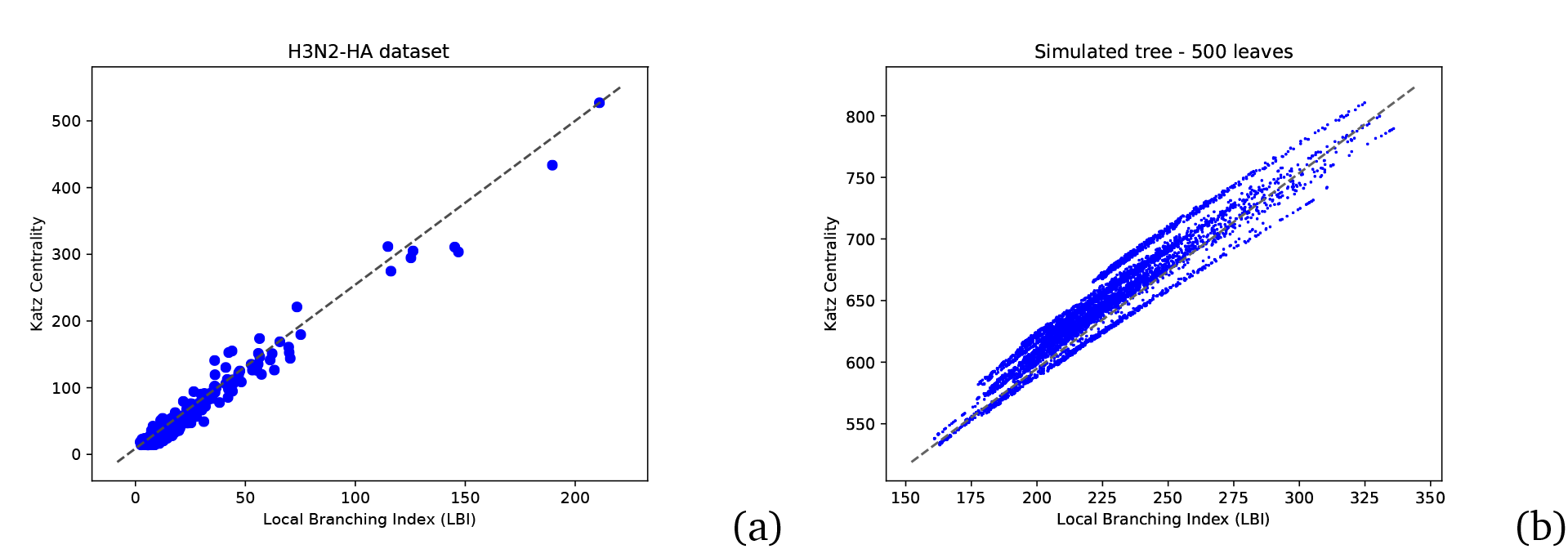
Correlation between Local Branching Index (LBI) and Katz Centrality measures (weighted version, where the distance between two nodes is defined as the sum of the branch lengths in the path separating them). (a) Each point corresponds to the LBI and Katz Centrality measures computed for the root of the 391 subtrees from the H3N2-HA dataset. Parameters used: *τ* = 50 (LBI), *α* = 0.95 (Katz Centrality). (b) Correlation for 10 trees simulated with a pure birth process. All trees have 500 leaves, and each point corresponds to the LBI and Katz Centrality measures computed for a node of those trees. Parameters used: *τ* =10 (LBI), *α* = 0.95 (Katz Centrality).

### Influenza virus B

We applied our method to influenza virus B. We used the same approach as for H3N2 and H1N1 to reconstruct the influenza virus B tree using sequences from 1980 to 2018-05 (for further details see Materials and Methods). The results of different experiments on influenza virus B are shown in Figure S9.

**Figure S9:**
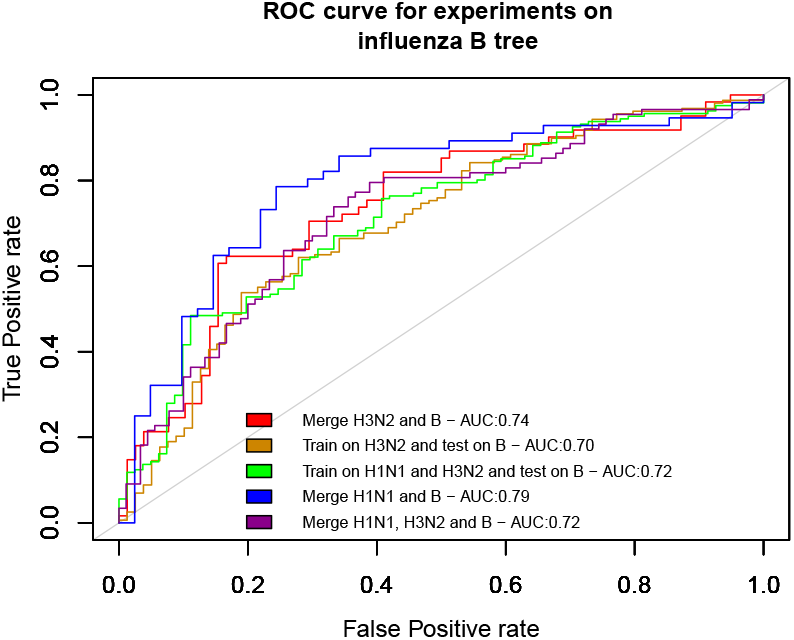
We find that a choice of *α* = 1.2 results in a good balance between successful and unsuccessful subtrees. Combining H1N1 and B subtrees results in good performance, suggesting similarities between the link between subtree structure and success between H1N1 and influenza virus B.

### Best Features

For robust feature selection we followed the ensemble technique introduced in [42]. They show that combining multiple (unstable) feature selectors yields more robust feature selection than using a single selection method. We use 4 models including logistic regression, random forests, SVM with linear kernel and learning vector quantization (LVQ)[19] to rank the features based on their contribution in the classification task. In the general classification of the H3N2 subtrees, the epitope features are among the most important. However, classification based only on epitope features reduces the AUC from 0.89 to 0.72, and our classifiers perform well on H1N1 (0.86 AUC, and 0.76 average AUC in the 10-fold cross validation) despite not having the epitope features. No single feature or small group of features that we have identified can perform as well as the combined phylogenetic and epitope features. We did not attempt to reduce the feature set to obtain a minimal set of features with the optimal performance.

### Predictions for the growth rate of WHO clades in 2019

**Figure S10:**
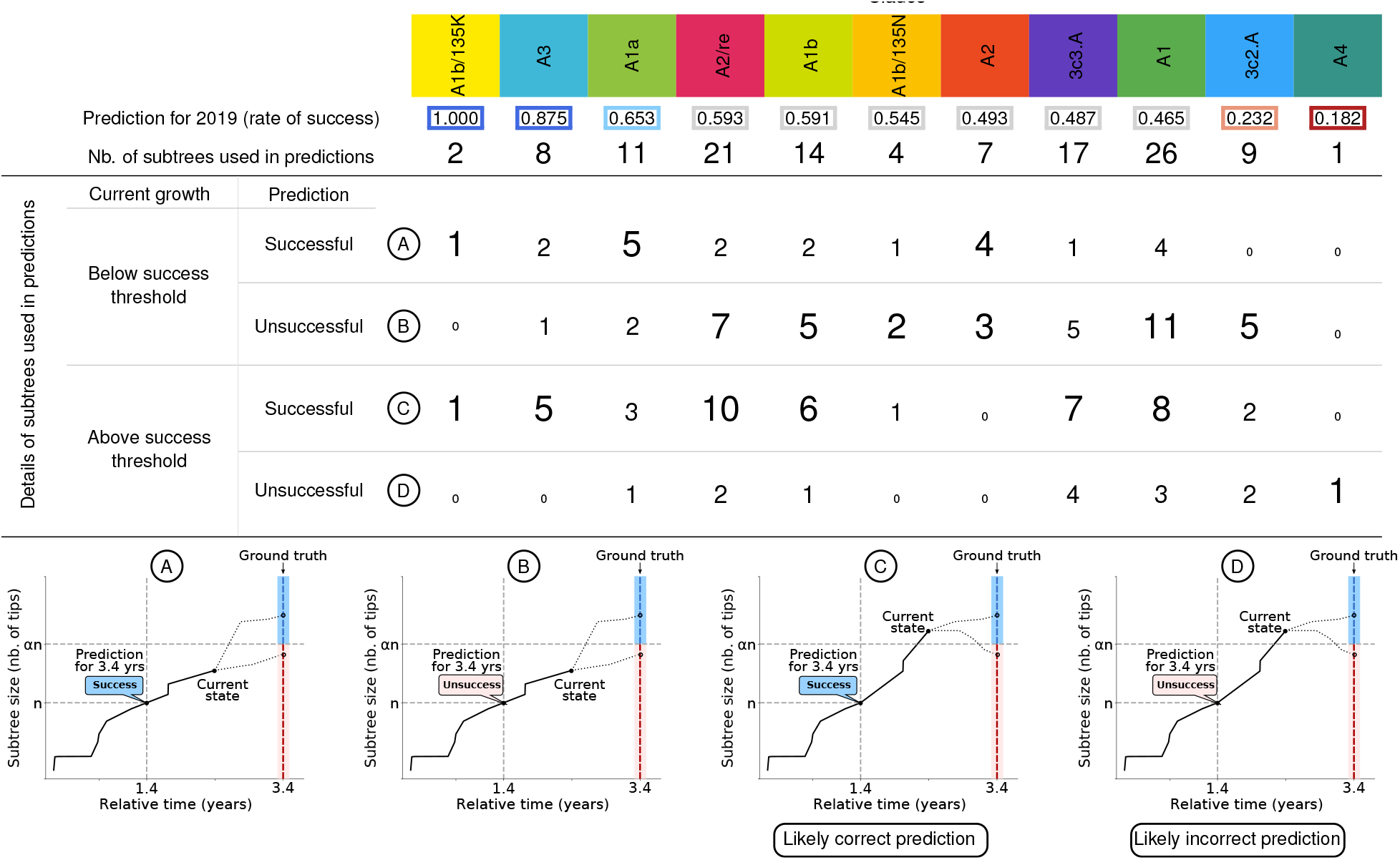
Predictions of the rate of success for the clades defined by the World Health Organisation (WHO). A column contains the information about one clade, indicated on the top and following the same color code as Figure 3. The columns (clades) are sorted according to the rate of success, from the most successful (left) to the least successful (right). The rate of success of a clade is an average of all predictions from subtrees whose majority of tips contain DNA markers linked to the clade (11 predictions per tree). All subtrees used in these predictions were extracted between 2015 and 2018 and, thus, have an unknown outcome, as the actual growth rate in 3.4 years from the initial node is still inaccessible. The subtrees are classified in four types: (A) subtrees whose prediction is to succeed but the current growth is still below the threshold considered to be successful; (B) subtrees whose prediction is to fail and the current growth is below the threshold considered to be successful; (C) subtrees whose prediction is to succeed and the current growth is above the threshold considered to be successful (notice that even in this scenario it is not possible to be certain, since the structure of the tree may change); (D) subtrees whose prediction is to fail but the current growth is above the threshold considered to be successful. Schematics for each case are presented on the bottom of the table.

